# Improved online event detection and differentiation by a simple gradient-based nonlinear transformation: Implications for the biomedical signal and image analysis

**DOI:** 10.1101/2020.08.16.253435

**Authors:** Anastasia Sokolova, Yuri Uljanitski, Airat R. Kayumov, Mikhail I Bogachev

## Abstract

Despite recent success in advanced signal analysis technologies, simple and universal methods are still of interest in a variety of applications. Wearable devices including biomedical monitoring and diagnostic systems suitable for long-term operation are prominent examples, where simple online signal analysis and early event detection algorithms are required. Here we suggest a simple and universal approach to the online detection of events represented by abrupt bursts in long-term observational data series. We show that simple gradient-based transformations obtained as a product of the signal and its derivative lead to the improved accuracy of the online detection of any significant bursts in the observational data series irrespective of their particular shapes. We provide explicit analytical expressions characterizing the performance of the suggested approach in comparison with the conventional solutions optimized for particular theoretical scenarios and widely utilized in various signal analysis applications. Moreover, we estimate the accuracy of the gradient-based approach in the exact positioning of single ECG cycles, where it outperforms the conventional Pan-Tompkins algorithm in its original formulation, while exhibiting comparable detection effectiveness. Finally, we show that our approach is also applicable to the comparative analysis of lanes in electrophoretic gel images widely used in life sciences and molecular diagnostics like restriction fragment length polymorphism (RFLP) and variable number tandem repeats (VNTR) methods. A simple software tool for the semi-automated electrophoretic gel image analysis based on the proposed gradient based methodology is freely available online at https://bitbucket.org/rogex/sds-page-image-analyzer/downloads/.

## 1. Introduction

Detection and differentiation of events is a long studied problem in signal analysis. While conventional solutions focused mainly on the calculation of simple statistics in a gliding window and simple threshold based detection rules, more recent approaches involve various sophisticated mathematical tools such as fluctuation analysis, pattern recognition, machine learning, empirical model decomposition and others [1–5]. Despite of the recent advancements in modern signal analysis methodology, leading to enhanced accuracy and performance, complex methods typically require considerable computational resources, as well as sufficient knowledge about specific signal properties (sometimes obtained during an inevitable preliminary learning procedure), that strongly limits their application. Wearable systems for biomedical diagnostics that are often required to operate on a long-term basis from days to months and even years without any technical intervention appear a prominent example [6]. Although long-term health monitoring systems have been successfully developed for decades, a more recent tendency is to facilitate them with certain emergency alert functionality that, in turn, requires certain preliminary signal analysis to be performed online [7]. Since long term operation requirements strongly limit the power supply and thus also the computational resources available for the online analysis, it is often based on rather simple signal processing algorithms of the past decades. While being clearly outperformed by their modern counterparts in the presence of excessive resources, they are nevertheless more suitable under limited resource conditions and still occupy this niche [8–10].

Long-term ECG monitoring systems appear prominent examples of wearable systems where various modifications of the Pan-Tompkins algorithm dating back to 1985 remain rather common solutions for the online detection of single ECG waveforms [11]. Since detection results are being used in further analysis, including differentiation of QRS shapes for cardiovascular diagnostics, their accuracy also affects the diagnostic results. A common problem in multi-lead ECG analysis is that the QRS shapes vary considerably between different leads as well as exhibit pronounced alterations in response to body postural changes and physical activity. As a result, particular QRS shapes can hardly be determined in advance, and thus using algorithms that are robust against shape variations remains a more suitable approach. Similarly, in molecular diagnostics protein mass spectra obtained under non-identical experimental settings vary considerably, since the variety of factors influencing the experiment outcome can hardly be reproduced even in high-tech labs.

In this paper, in order to overcome typical limitations of existing methods, we suggest using a simple gradient-based transformation obtained as a product of the signal and its derivative prior to running the detection algorithm. As we show below using both synthetic and empirical data analysis, the proposed transformation leads to the improved accuracy online detection of any significant bursts in the observational data series irrespective of their particular shapes.

## 2. Materials and Methods

### 2.1. Data sources

The first class of data used in our analysis includes 48 half-an-hour long two-lead ECG recordings of 24 male aged 32 to 89, and 23 female subjects aged 23 to 89, referred to the Arrhythmia Laboratory at Boston’s Beth Israel Hospital (now the Beth Israel Deaconess Medical Center) obtained from the MIT-BIH Arrhythmia Database; 72 half-an-hour long twelve-lead ECG recordings of 17 male and 15 female subjects aged 18 to 80, referred to the St. Petersburg Institute of Cardiological Technics (Incart), St. Petersburg, Russia; as well as 33 fragments preselected from 18 long-term ECG recordings from 5 male subjects aged 26 to 45, and 13 female subjects aged 20 to 50 from the MIT-BIH Normal Sinus Rhythm Database. All recordings have been obtained from the PhysioNet databank [12]. Neither of the recordings contained any significant arrhythmias.

The second class of data includes protein mass spectra obtained by Polyacrylamide Gel Electrophoresis (SDS-PAGE) in an earlier work [13]. The total protein was measured by Bradford assay using BSA for standard curves. Normalized samples containing 5 µg of total protein per lane were separated in 10% – 15% (w/v) separating polyacrylamide gels with 4% (w/v) stacking gels under denaturing conditions, as described in [14]. The gels were stained with either Coomassie Brilliant Blue or silver nitrate, as described in [15], and documented on trans-white mode.

The final images contain lanes with intensity that is proportional to the concentration of proteins with specific mass. Therefore, by considering the intensity as a function of the coordinate along the studied lane, one obtains a functional curve that is proportional to the protein concentration as a function of the logarithm of the protein mass.

### 2.2. Event detection methods

Several rather universal methods for event detection are well established in conventional signal theory, and represent the basis for multiple common algorithms utilized in various signal analysis applications. In our work, we consider the following algorithms, as well as their modifications and/or combinations:

#### The correlation algorithm

is based on the calculation of the variants of the inner product 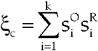, such as the cosine similarity, Otsuka-Ochiai coefficient [16, 17], or correlation coefficient between the observational signal samples 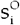 taken in a gliding window of *k* samples and a reference shape 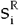 of the same size. Although the correlation algorithm is known as a theoretically optimal solution under rather general assumptions, it requires preliminary knowledge of the shape of the reference signal s^R^. In practical scenarios, the QRS shape can either be determined from a preliminary study of the same patient under more or less similar conditions, or estimated from the previously observed signal fragment by averaging with elimination of anomalous measurements in order to reduce the noise influence [18]. Neither of the above scenarios takes into account rapid variations in QRS shapes that strongly limit the application of this otherwise highly accurate approach.

#### The energy based detection algorithm

focuses on the calculation of the local observational signal energy by integration of the squared signal 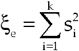 in a gliding window of *k* samples. While this algorithm represents one of the simplest possible solutions with a minimum number of free parameters, one has to take into account that the signal energy varies considerably between different ECG leads and is further affected by postural changes. Although the threshold value for a given window size can be adjusted using some adaptive algorithms [19], this adaptation often appears insufficient to take into account pronounced shape alterations. Moreover, in anomalous ECG records the QRS can be inadvertently confused with other ECG waves such as P-wave and/or T-wave leading to inaccurate heartbeat timing assessment.

#### Pan-Tompkins algorithm

in its original formulation is based on the analysis of the squared derivative 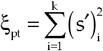 of the raw signal, smoothed over five consecutive raw signal samples to reduce the impact of the high-frequency noise, and aggregated in a time window of *k* samples [11]. Since this algorithm relies on the values of the derivative, it successfully detects any abrupt changes in the signal, while remaining barely sensitive to the particular signal shape analyzed. Due to its simplicity and universality, different variants of the Pan-Tompkins algorithm remain a standard solution for wearable ECG analysis systems over several decades [20]. The disadvantage is that the derivative remains quite sensitive to the high-frequency noise components mainly represented by electrophysiological activity of the skeletal muscles, especially in those leads where the ECG signal amplitude (and thus also energy) are small. In addition, abrupt ECG changes are characteristic for the artifacts induced by temporary detachment or shift of the electrodes, that typically occurs due to body motion during physical activity (e.g. exercise). Of note, using smoothed derivative estimates partially enhances its robustness against high-frequency noise, at the cost of reduced temporal resolution.

#### The modified energy based detection algorithm

that implies the analysis of a weighted linear combination of the raw signal and its derivative 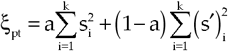 has been suggested recently [21]. Although in this recent study it has been demonstrated that improved performance could be achieved by combining the signal energy based and the derivative based analysis compared to using only one of them, the choice of the optimal weighting coefficients remains challenging, as it strongly depends on particular signal shapes, and thus also varies between individuals, leads, time fragments and so on. In turn, these variations require additional adjustments of the weighting coefficient, this way increasing the complexity of the algorithm [21].

#### The gradient based algorithm

that we propose here focuses on a similar analysis of the local signal gradient estimate obtained as the product of the raw signal and its derivative 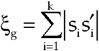, this way providing a kind of superposition of the energy based detection and Pan-Tompkins algorithm. As we show below, while remaining sensitive to any events characterized by rapid changes in the local signal energy, the gradient based algorithm appears robust against signal shape alterations, while requiring no additional free parameters.

In practical scenarios, the exact signal shape *s*_*i*_ can never be determined, although it could be estimated from its noisy observations *x*_*i*_. Let us assume that *x*_*i*_ *= s*_*i*_ *+ n*_*i*_, where *s*_*i*_ is the exact (noise-free) signal shape, and *n*_*i*_ is the additive background noise. As a result, in the above statistics ξ that are used for decision making, signal samples *s*_*i*_ are being replaced by noisy observations *x*_*i*_, leading to inevitable errors. In the ECG records, one can usually estimate *n*_*i*_ at least indirectly by subtracting an approximation of *s*_*i*_ from *x*_*i*_ in a gliding window. Under the assumption that the noise samples *n*_*i*_ are independent and identically distributed with zero mean, the simplest and most straightforward way to approximate *s*_*i*_ could be obtained as a moving average of the raw signal samples *x*_*i*_. In a more general scenario, a polynomial approximation, typically either linear or parabolic, is commonly used [22]. In the SDS-page fingerprinting, *n*_*i*_ can be determined directly by considering an empty lane, where no useful experimental sample has been added, while the other effects ranging from chemical noise to optical effects are generally similar to other lanes containing signals of interest.

### 2.3. Pattern differentiation methods

For best distinguishing between the studied signal shapes (in our case exemplified either by the QRS waves or by the characteristic protein mass spectra bands), we focus on a technique based on the basic decision theory principles. Let us assume that the observational signal 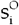 is compared against *M* possible reference signals 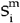, where m=1,2,…,*M*, and that the reference signal 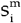 that best corresponds to the observational signal 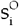 should be found.

In decision theory, assuming that *n*_*i*_ are Gaussian distributed, the optimal similarity metrics are based on the normalized inner product 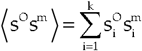 known as the cosine similarity score

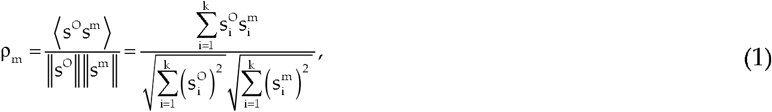

that is bounded between −1 and 1, or 0 and 1 for non-negative signals (like the image intensity). Since we have no access to the exact shapes of 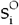 and 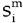, due to the influence of multiple random factors embedded in the noise samples *n*_*i*_, we have to rely upon estimations of the relevant similarity score

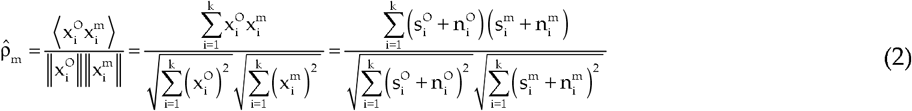

from measured 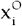 and 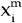, which, in turn, are random variables characterized by its own distribution 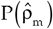 [19]. In most practical scenarios, there is certain overlap between the distributions 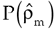 for different reference signals *m1* and *m2* corresponding to different tested hypotheses, i.e., for the similarity scores between 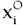 and 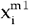 vs 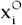 and 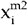, respectively. This overlap leads to inevitable misclassifications when signal *m1* is recognized as signal *m2* and vice versa, constituting the so-called confusion matrix. Accordingly, best differentiation can be achieved between reference shapes 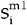 and 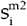 with smallest cross-correlation ρ = −1, exemplified by black & white lanes in inventory barcode systems.

## 3. Validation of the proposed methodology: an analytical treatment

### 3.1. Event detection strategy

In the following, we consider an analytical approach to the comparison of different event detection methods. Although the correlation analysis based approach provides with the optimized solution under rather general conditions, it requires the preliminary knowledge of the reference shape. In biomedical signal analysis, the above condition can hardly be met, as there are pronounced discrepancies between signal shapes observed in different individuals, depending on their condition, and on the registration method. In addition, pronounced variability of the analyzed signal shapes strongly limits the usability of the same reference shape over entire long-term recordings.

For the above reasons, we do not consider the correlation based approach to the event detection as a relevant tool in our case, although in some tests we use it as a reference. Using this comparison, one may consider to which extent the *a priori* knowledge of the signal shape could potentially improve the results.

Let us assume that the signal *s*_*i*_ is hindered by measurement noise *n*_*i*_ leading to the observation *x*_*i*_ *= s*_*i*_ *+ n*_*i*_. For appropriate comparison of all methods, we consider the same algorithm design based on the calculation of statistics ξ in a gliding window of *k* samples followed by decision making based on the comparison of ξ against a decision threshold Θ. To characterize the noise level, we use the signal-to-noise ratio 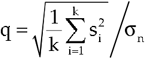. Since the noise is induced by multiple independent sources, let us also assume that *n*_*i*_, at least in the first approximation, is represented by Gaussian white noise with zero mean and variance 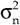. Under this scenario, for an arbitrary signal shape 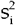 one expects also Gaussian distribution of the statistics ξ used for the decision making [19]. Accordingly, the probability density function (PDF) of the statistics ξ used for decision making can be written as

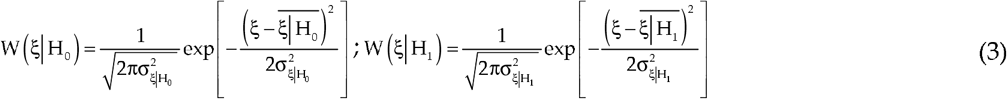

both for the null hypothesis corresponding to the absence of event leading to *x*_*i*_ *= n*_*i*_ and for the alternative hypothesis corresponding to the presence of event *x*_*i*_ *= s*_*i*_ *+ n*_*i*_, respectively. When the statistics ξ is compared against a certain threshold Θ, the decision in favor of hypothesis H_1_ made if ξ <Θ, and decision in the favor of hypothesis H_1_ is made if ξ >Θ. The probability that the observational statistics satisfies the latter condition can be expressed as

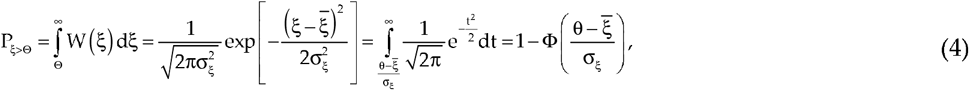

where 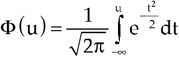 is the cumulative distribution function of the standard Gaussian distribution. In turn, Equation (4) is known as the complementary cumulative distribution function (CCDF), or exceedance probability, a quantity widely used in classification analysis [23]. In terms of exceedance probabilities, the accuracy of the decision making is determined by the true positive rate indicating how often the decision in favor of the hypothesis H_1_ is made when the hypothesis H_1_ is actually valid, and by the false positive rate, also known as Type I error rate, indicating how often the decision in favor of the hypothesis H_1_ is made when in fact the hypothesis H_0_ is valid. The above quantities can be obtained directly from the CCDFs by substituting the unconditional ξ by the conditional ξ |H_1_ and ξ| H_0_, respectively. The corresponding procedure leading to the receiver operator characteristic (ROC) curve [24] quantifying the efficacy of the algorithm and the error rates is illustrated in Fig. 1.

**Figure 1.**
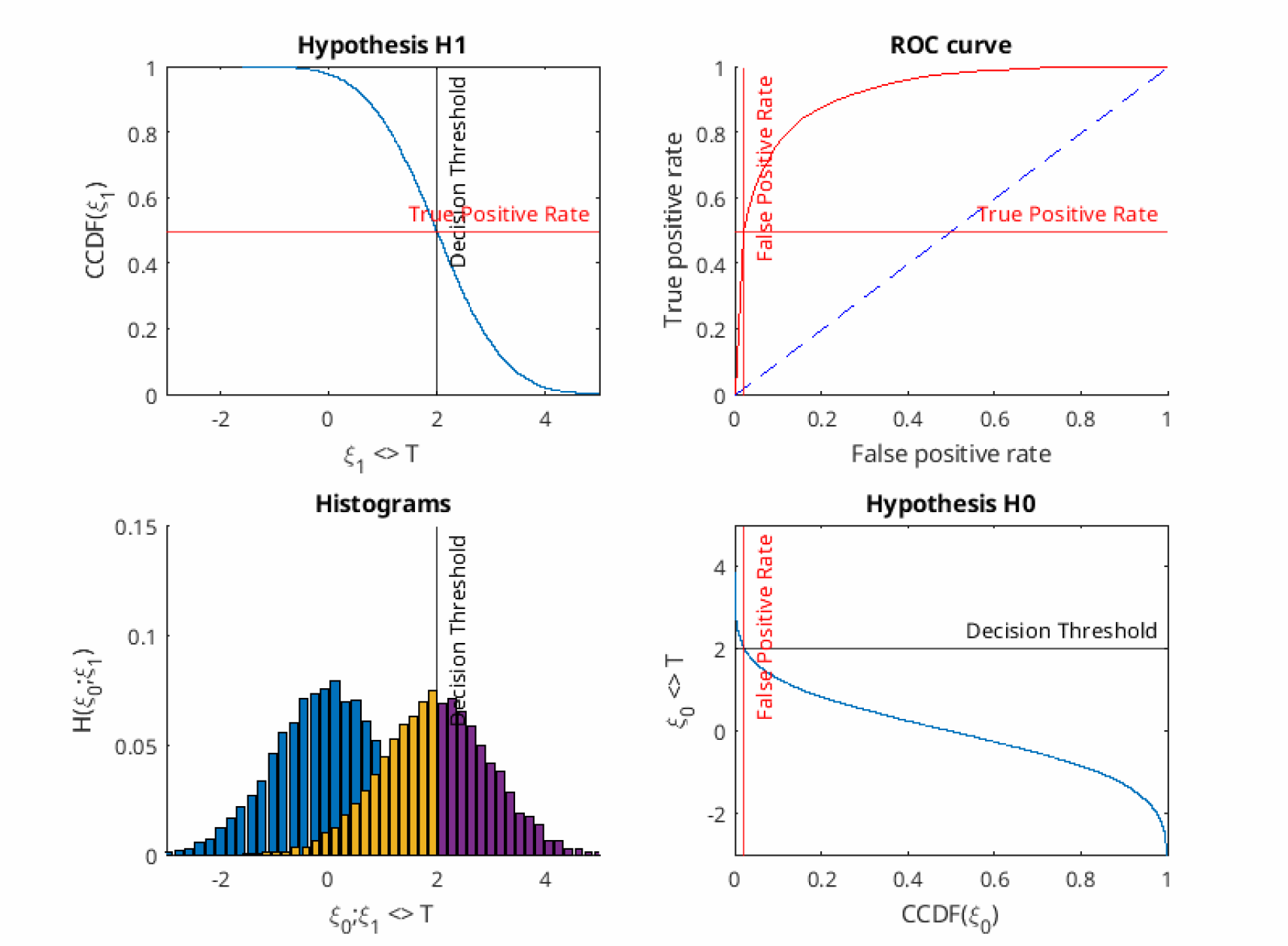
Illustration of the ROC-curve estimation.

In turn, as the distributions W (ξ) are Gaussian, they are fully determined by only two parameters, the average value 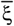 and the standard deviation σξ. Accordingly, to obtain the full functional form of the CCDFs, it is also sufficient to know only the averages and the standard deviations. In the following, the above quantities are being estimated analytically for each of the decision statistics.

**In** the **energy based detection algorithm**, 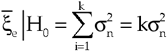 and 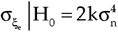, while 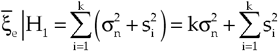 and 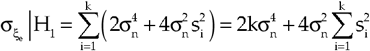.

**In the Pan-Tompkins algorithm**, the assumption that the smoothed derivative is based on a 5-point estimate given by 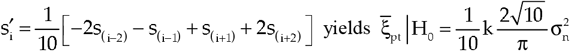

and 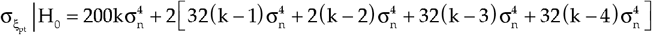, while 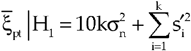 and 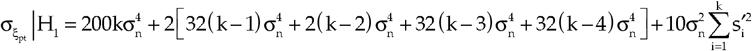.

**In the gradient based algorithm**, given that 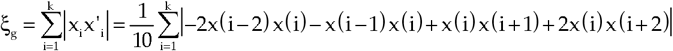

similar transformations lead to 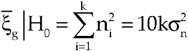 and 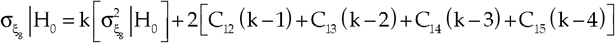, while 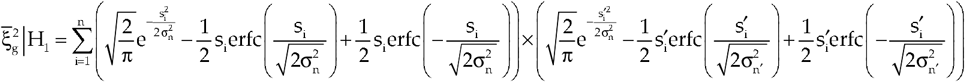 and 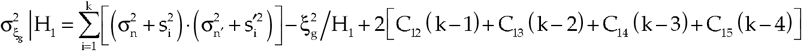

where 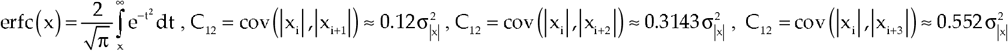 and 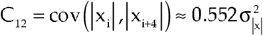 are the cross-correlation coefficients between the samples within the smoothed derivative estimation window.

### 3.2. Signal shape differentiation strategy

As already noted above, for the best differentiation, two signal shapes should be characterized by the lowest possible similarity score, ideally approaching −1 for the pairwise comparison scenario, with optimized design strategies leading to ensembles of *M* signals characterized by ρ _*ij*_= −1 (M − 1) or ρ_*ij*_ = 0 for the suboptimal but easier synthesized orthogonal signal ensembles [22].

However, for the majority of natural observational signals, the situation is nearly the opposite, i.e., different ECG waves, normal and pathological QRS shapes exhibit only moderate (although characteristic) differences, and proteomes of taxonomically close species as well as treated and untreated cultures typically differ by few proteins only, and thus their similarity scores typically approach one. In this case, the errors are inevitable if the above methodology is applied straightforwardly, due to the overlap of the corresponding distributions, see Fig. 1. The simplest way to reduce the overlap is to remove the common part from all references that can be achieved by subtracting the average of all references in a given set from each of them leading to broader distribution of 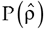 between −1 and 1. Other alternatives include different nonlinear transformations of the raw signals.

Next there are two ways how to proceed with decision making. The first scenario assumes that the tested signal is believed to have a proper reference in the considered reference set. In this case, the solution is straightforward and based on the selection of the reference with the highest similarity score with the studied sample. The second scenario assumes that there is no certainty, whether a proper reference can be found in the considered reference set. In this case, with at least an approximate knowledge of 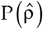, the decision threshold Θ can be set at a certain (e.g. 5%) quintile of 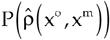. Exceedance of this threshold by the estimated 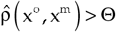 indicates that with 95% confidence probability there are no significant differences between the compared signals x° and x^m^. Of note, this second scenario includes potential ambiguity, leading to possible indications that (i) neither of the reference samples correspond to the test sample with 95% confidence probability, or (ii) there are several signals x^m^ m=1,2,…,r satisfying 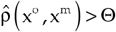 indicating that neither of the r reference signals exhibit statistically significant differences with the test signal x^m^.

In practice, 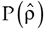 can be obtained both numerically and analytically. The numerical estimation is straightforward and is based on Monte-Carlo simulations following (2) with different noise samples obtained e.g. from an empty lane. Analytical treatment that does not require any iterative computations can be considerably simplified under the assumption that 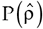 is Gaussian distributed, and thus can be fully determined by its mean 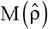 and variance 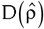 [19]. In this case,

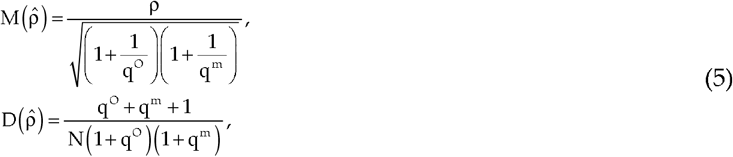

where ρ is the cosine similarity score between s° and s^m^, while

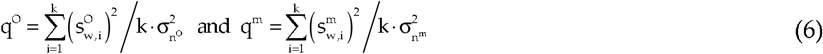

are the signal-to-noise ratios characterizing the noisiness of the sample and the reference signals, while 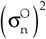 and 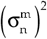 are the noise variances for the observational and the *m*-th reference signals, respectively. The noise influence can be further reduced by signal averaging, for example, in the SDS-PAGE electrophoresis gel image analysis each sample in the resulting signal can be obtained by averaging over the lane width w [25–27]. In this case, under the assumption that the image is hindered by centered white noise, the averaging procedure leads to the enhancement of the effective signal-to-noise ratio by a factor of w

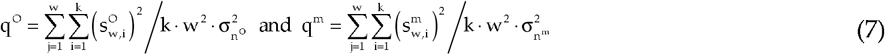

where w° and w^m^ are the observational and the *m*-th reference lanes widths, as well as 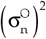 and 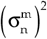 are the noise variances for the observational and the *m*-th reference signals, respectively, and k is the lane length in pixels. An important consequence of (7) is that taking wider signal lanes leads to the reduction of the effective noise levels due to noise averaging, in agreement with known results from signal theory, and thus is preferential. We also like to note that the above methodology appropriately accounts only for the instrumental noise, while does not take into account other random factors such as e.g. inter-specimen variability. To account for the latter, reference lanes could be merged from experiments obtained for several specimen.

## 4. Results and Discussion

### 4.1. Detection of QRS waves in long-term ECG records

In the following, we further validate the proposed approach by comparing analytical results with estimates obtained in a numerical treatment, and next apply them to the QRS detection in the context of an automated long-term QRS analysis problem.

In the QRS detection problem, we assume that the signal energy is mostly concentrated in the R-wave. Accordingly, in the first approximation, the corresponding signal can be represented by a simple triangle-shaped model depicted in Fig. 2a. The methodology described above have been reproduced in a numerical simulation, and compared to the results of the analytical treatment. The ROC-curves exemplified in Fig. 2b for q=0.1 indicate a perfect match. Next by fixing the false positive rate (corresponding to a vertical cutoff in Fig. 2b), the noise robustness characteristics (NRC) were obtained by finding the crossing points of this vertical cutoff with the corresponding ROC-curves [28]. Fig. 2c shows the NRC-curves exemplified for the fixed false positive rate α=0.1.

**Figure 2.**
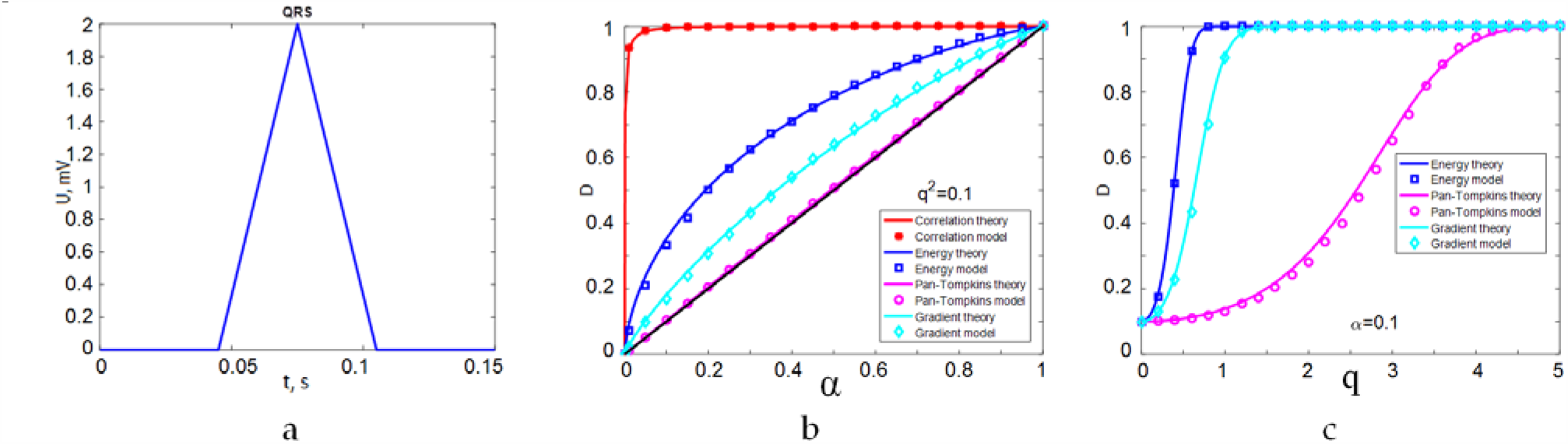
(a) A simplified model of the QRS shape; (b) ROC-curves and (c) NRC-curves characterizing the efficacy of the QRS shape detection.

The figure shows that, as expected, the correlation based algorithm yields the best efficacy, although its application requires the *a priori* knowledge of the particular QRS shape. Since the above requirement makes this approach very impractical due to the QRS shape variability, it is skipped in further analysis. As indicated by the ROC-curves and by the NRC-curves for the remaining algorithms, the energy based one is the second best, although the gradient based algorithm appears only slightly less efficient. In contrast, the Pan-Tompkins derivative based algorithm is characterized by considerably lower true positive rates D for a broad range of signal-to-noise ratios q=0.5…4.

Next we consider the probability of the confusion between the QRS wave and other ECG waves such as P-wave and/or T-wave. In contrast to the typically peak-shaped QRS, both P- and T-waves are characterized by rather smooth shapes that could be in the first approximation modeled by a half-period of the cosine function. To a reasonable approximation, we next assume that the energies of the two different waves under comparison are equal.

Fig. 3a shows the corresponding signal shapes, while Fig. 3b shows their derivatives. Finally, Fig. 3c shows the ROC-curves for the differentiation problem, now assuming that the hypothesis H_0_ corresponds to the presence of the half-period-cosine-shaped P- or T-wave, while the hypothesis H_1_ corresponds to the presence of the QRS wave. Our results indicate that the successful differentiation requires considerably higher signal-to-noise ratios. Again, the correlation based algorithm appears the most accurate, although for the reasons specified above we used it here as a reference method only. Among other algorithms, the Pan-Tompkins derivative based algorithm appears more efficient. However, the above results indicate that it is characterized by considerably lower noise robustness (see Fig. 2c). In contrast, the gradient based algorithm appears only moderately less efficient in both detection and differentiation scenarios (compare Figs. 2 and 3), making it a promising tool for the automated ECG analysis.

**Figure 3.**
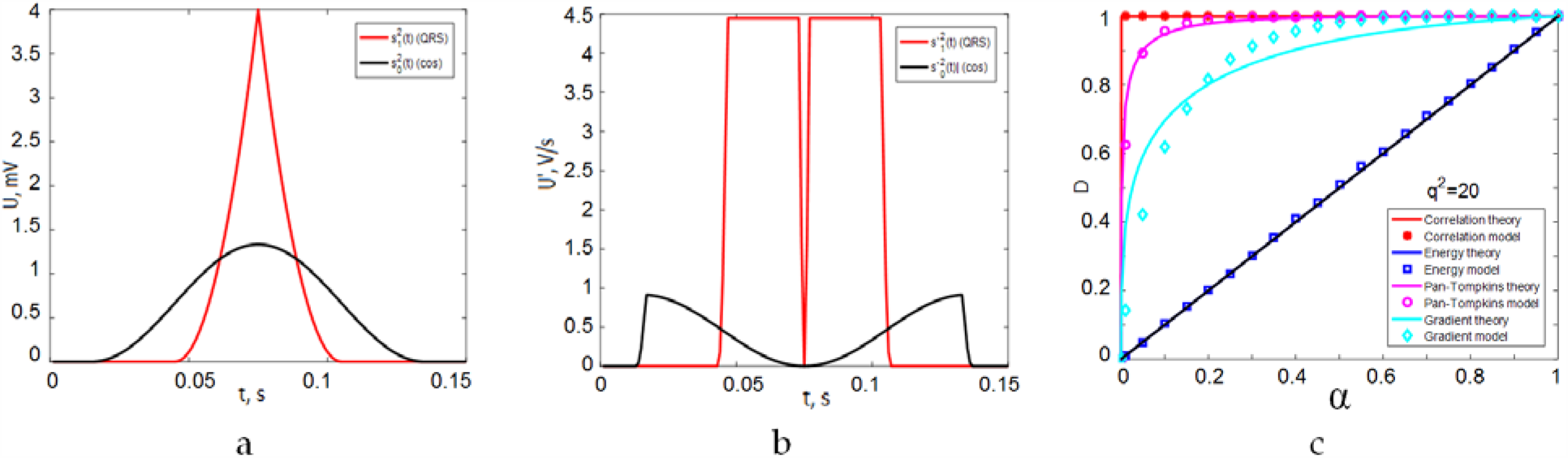
(a) Simplified models of the QRS-wave, as well as of the P- or T-waves; (b) the derivatives of the signals shown in Fig. 2a; (c) ROC-curves characterizing the efficacy of the QRS shape differentiation.

For the best performance of the corresponding algorithms, the appropriate window size *k* has to be chosen. Since the statistics used for decision making are averaged in a gliding window, the relative duration of the signal peak associated with the event of interest (such as the QRS waves in the ECG) as a fraction of the gliding window size is the key quantity that determines the detection results, while also affecting the ‘abruptness’ of the peak to be detected. In a theoretical setting, it is rather evident, that with the increase of the window size, the relative enhancement of the local maximum of the decision statistics would increase as long as the gliding window is below the QRS size, maximize when in becomes equal to the QRS size, and decrease when window size exceeds the QRS size. Thus the optimal window size should be equal to the QRS size for the optimized detection.

However, in most practical scenarios, the duration of the QRS shape varies considerably with varying heart rate, conditions, and with shape alteration associated with various anomalies, such as some types of arrhythmia. Although normal QRS shapes are typically of 0.06-0.1s duration, QRS elongation above 0.12s can be observed in the presence of ventricular extrasystoles, that requires elongation of the gliding window. However, for longer windows there is an increased risk that the other ECG waves outside the QRS, such as P- and/or T-waves, may be included into the window simultaneously with the QRS. Thus optimization of the window size k is required prior to further application of the corresponding methods and algorithms.

In the following, we proceed with the empirical data analysis. Fig. 4 shows the false positive rate (FP), the false negative rate (FN), and the total error rate (FP+FN) as a function of the gliding window size *k* obtained for the MIT-BIH Normal Sinus Rhythm Database. The figure shows that for all algorithms, the lowest total error rate can be observed around 0.18±0.01 s gliding window duration. As a side remark, we also like to note that the Pan-Tompkins algorithm appears more sensitive to the accurate selection of the gliding window duration, indicated by a V-shaped total error rate around the minimum observed at *k* = 0.19 s, while the other two algorithms appear less sensitive, characterized by rather U-shaped total error rate curves, that could be their additional advantage, since the accurate optimization of window size *k* could hardly be performed based on *a priori* information.

**Figure 4.**
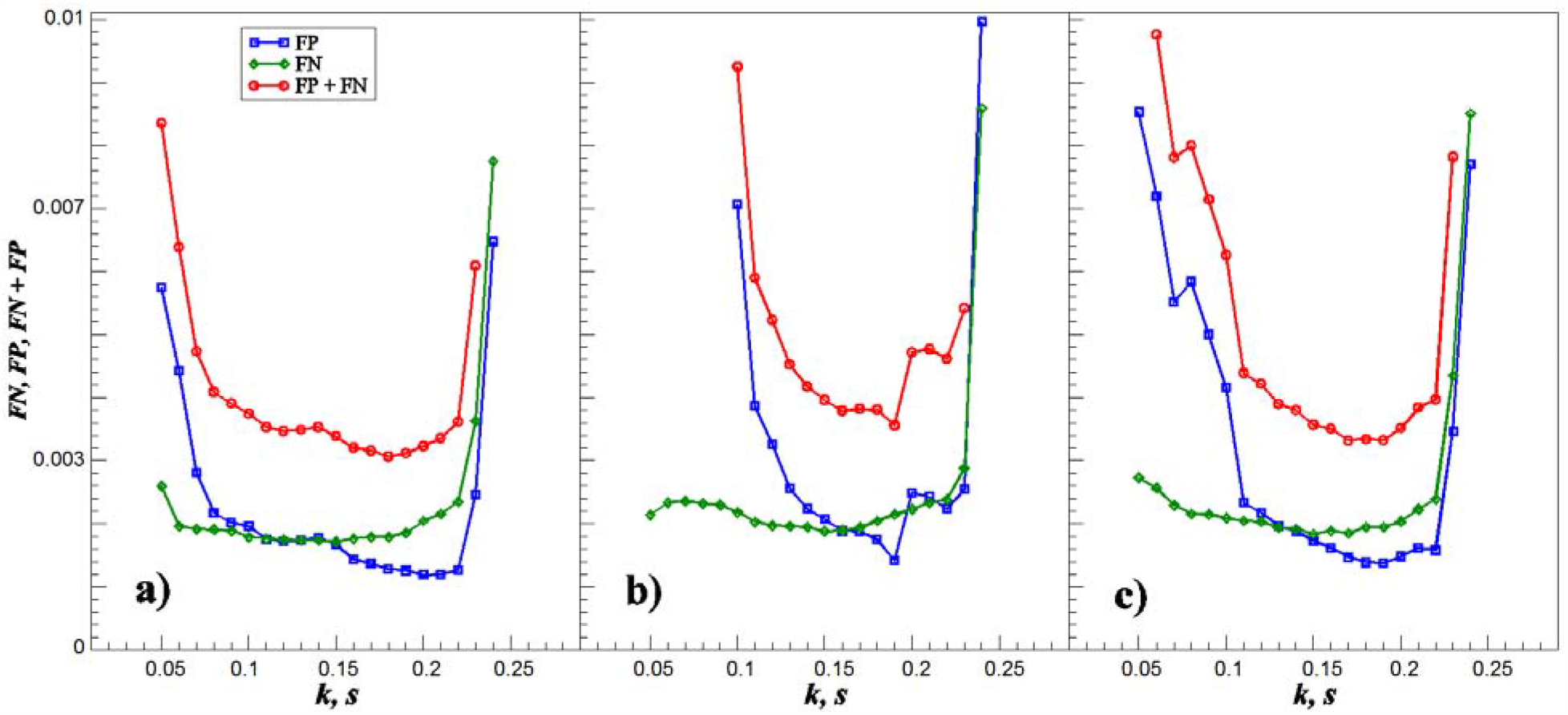
False positive rate (FP), false negative rate (FN), and the total error rate (FP+FN) as a function of the gliding window size *k* (in seconds) achieved by the (a) energy based, (b) Pan-Tompkins, and (c) gradient based algorithms for the MIT-BIH Normal Sinus Rhythm Database.

Fig. 5a exemplifies an ECG record fragment containing significant QRS shape variability, including ventricular arrhythmic heartbeats also obtained from the PhysioNet databank. Figs. 5b-e show the derived statistics ξ used in the correlation, in the energy based, in the Pan-Tompkins and in the gradient based algorithms, respectively. Vertical lines in Fig. 5a denote the QRS position points verified by expert assessment that can be used as reference, while vertical lines in Figs. 5b-e correspond to the points where QRS waves have been detected by the corresponding algorithms.

**Figure 5.**
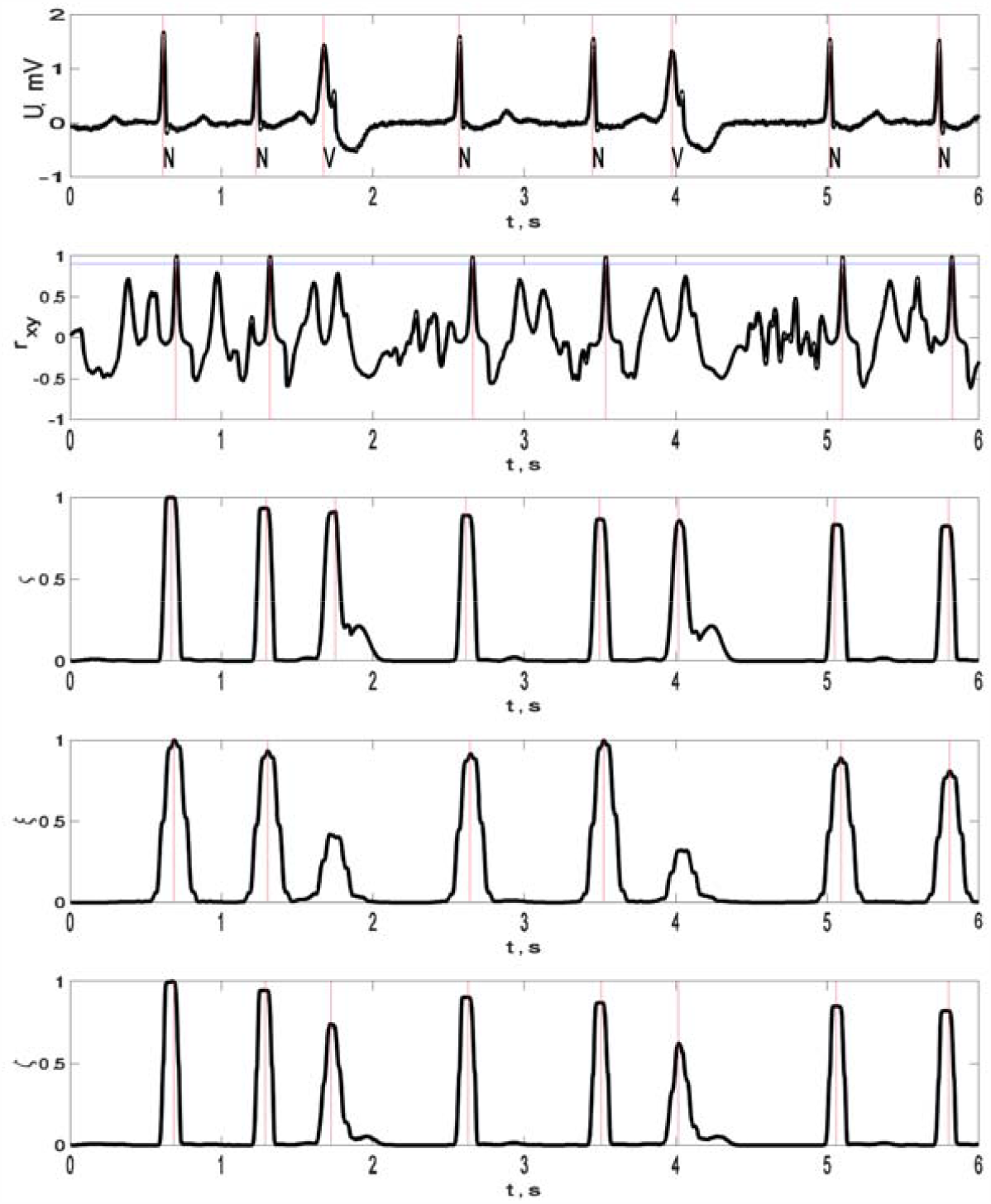
(a) Example of an ECG **record** fragment; the derived statistics ξ used in (b) the correlation based, (c) the energy based, (d) Pan-Tompkins and (e) the gradient based algorithms. Vertical lines denote QRS positions determined by (a) an expert assessment as well as by (b)-(e) the corresponding automated detection algorithms.

Finally, we tested how the algorithms performed being applied to the observational long-term ECG recordings. First, we have analyzed the efficacy of QRS detection for the 48 recordings from the 2-lead ADB database and 72 recordings from the 12-lead Incart ECG database indicated by the corresponding F1-scores. Figs. 6 and 7 show boxplots and p-values obtained using the Mann-Whitney U-test for the respective datasets. The efficacy has been quantified by comparing against the manual reference obtained from the corresponding databases. The figure shows that in both cases approximately one-half of all ECG leads appear suitable for an accurate QRS detection. Moreover, the overall detection efficacy can be improved by averaging the decision statistics over all leads. For the latter scenario, the efficacy is rather high indicated by F1-score median values around 0.9998 for the both databases. Moreover, there are no significant differences between the methods efficacy indicated by p > 0.05 (Mann-Whitney U-test).

**Figure 6.**
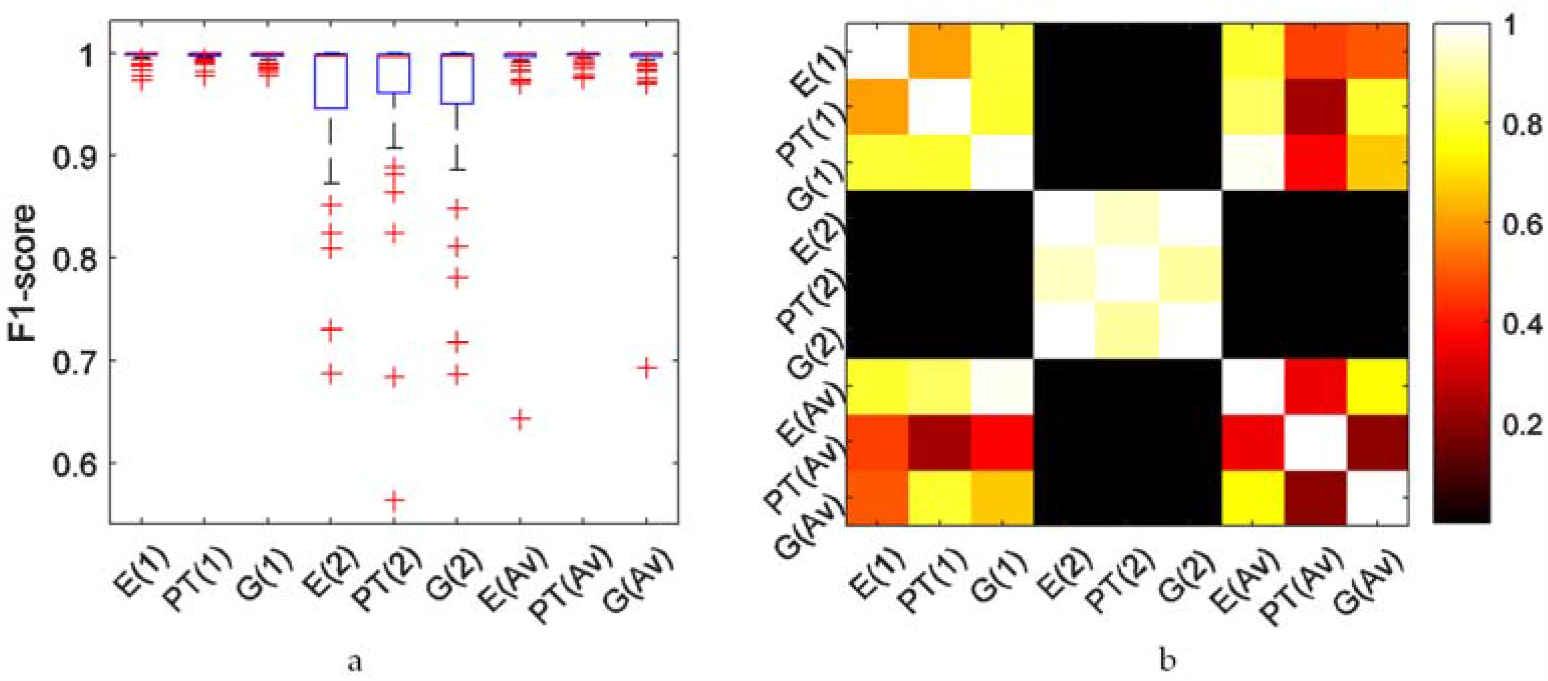
Boxplots. (a) and Mann-Whitney U-test p-values (b) for the F1-scores quantifying the efficacy of QRS detection for the 2-lead ADB ECG database. E(i), PT(i), and G(i) denote the signal energy based, Pan-Tompkins, and gradient based method, respectively, for the *i*-th ECG lead. The last three entries show similar results for the decision statistics averaged over all ECG leads.

**Figure 7.**
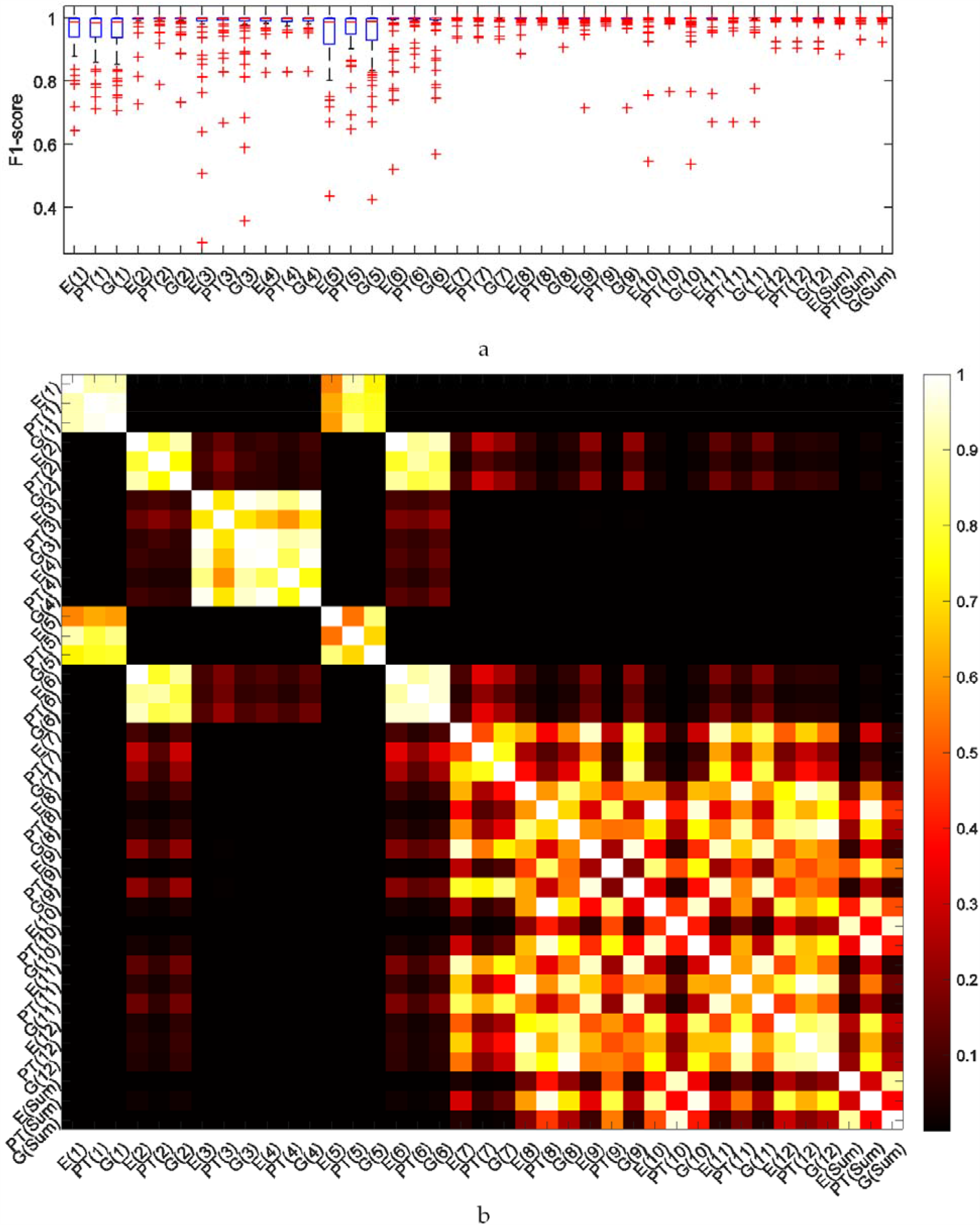
Boxplots. (a) and Mann-Whitney U-test p-values (b) for the F1-scores quantifying the efficacy of QRS detection for the 12-lead Incart ECG database. E(i), PT(i), and G(i) denote the signal energy based, Pan-Tompkins, and gradient based method, respectively, for the *i*-th ECG lead. The last three entries show similar results for the decision statistics averaged over all ECG leads.

Second, we have analyzed how accurately the exact position of the QRS shape is being fixed by either of the methods. For this test, 33 rather stationary fragments from the MIT-BIH Normal Sinus Rhythm Database have been chosen, to guarantee the ability of the correlation algorithm to detect QRS shapes. Of note, the above conditions are preferential for the correlation based algorithm, while the presence of arrhythmias typically reduces its accuracy, thus further confirming the requirement of the algorithms that are barely sensitive to the QRS shape.

A simple way would be to compare to the manual reference obtained from the database, but it might be also subjective in some cases. To overcome this limitation, we follow a more general and robust approach, based on the analysis of the total variance of the heartbeat intervals. Each ECG record is characterized by some natural heartbeat interval variability with variance 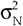, and some additional measurement variability characterized by variance 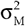. Under the assumption that these two sources of variability are independent, the total variance of the heartbeat intervals is given by the sum of the two variances 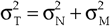. Although we do not know the exact 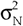 values, since it characterizes a natural process that is unaffected by the choice of the measurement algorithm, we can conclude that higher total variances 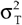 correspond to those algorithms that lead to higher measurement errors characterized by 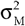. Following this strategy, we compared the total heartbeat interval variances 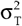 obtained by each of the algorithms tested. Since in earlier works preliminary filtering of the raw ECG signals have been suggested [21], although its impact on the results is twofold, resulting in the noise reduction accompanied by the inevitable loss of temporal resolution, we have tested the energy based and the gradient based algorithms both with and without preliminary filtering procedure with the use of a 4^th^ order Chebyshev filter. Corresponding results are summarized in Table 1.

**Table 1.** The total heartbeat interval variances 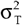 obtained by each of the algorithms for a series of observational long-term ECG records.

| Record No. | Pan-Tompkins | Energy based |  | Gradient based |  | Correlation based |
| --- | --- | --- | --- | --- | --- | --- |
|  |  | No filter | With filter | No filter | With filter |  |
| 1 | 0.05731 | 0.01334 | 0.04815 | 0.01593 | 0.03559 | 0.01268 |
| 2 | 0.06882 | 0.03240 | 0.05351 | 0.03241 | 0.04944 | 0.02655 |
| 3 | 0.03823 | 0.03282 | 0.03782 | 0.03246 | 0.04176 | 0.02701 |
| 4 | 0.06179 | 0.03094 | 0.06255 | 0.02849 | 0.05446 | 0.02584 |
| 5 | 0.04724 | 0.03193 | 0.05524 | 0.03287 | 0.04482 | 0.02839 |
| 6 | 0.03972 | 0.02504 | 0.03582 | 0.02327 | 0.03764 | 0.01761 |
| 7 | 0.04333 | 0.02342 | 0.03222 | 0.02096 | 0.03785 | 0.01789 |
| 8 | 0.06431 | 0.01367 | 0.03711 | 0.01602 | 0.02789 | 0.01233 |
| 9 | 0.05958 | 0.01778 | 0.04012 | 0.01571 | 0.03398 | 0.01245 |
| 10 | 0.06899 | 0.04343 | 0.07074 | 0.04389 | 0.06035 | 0.06602 |
| 11 | 0.05710 | 0.04352 | 0.06526 | 0.04327 | 0.05134 | 0.04372 |
| 12 | 0.09519 | 0.05070 | 0.09212 | 0.05180 | 0.06956 | 0.04819 |
| 13 | 0.09476 | 0.05066 | 0.11518 | 0.16829 | 0.10898 | 0.04816 |
| 14 | 0.07350 | 0.03022 | 0.07317 | 0.03127 | 0.04687 | 0.03039 |
| 15 | 0.06898 | 0.03514 | 0.07290 | 0.03270 | 0.04289 | 0.03060 |
| 16 | 0.04696 | 0.04301 | 0.04521 | 0.04325 | 0.04325 | 0.04110 |
| 17 | 0.06409 | 0.04179 | 0.06366 | 0.04257 | 0.05155 | 0.04080 |
| 18 | 0.05295 | 0.03633 | 0.04984 | 0.03591 | 0.05093 | 0.02950 |
| 19 | 0.02872 | 0.02938 | 0.02872 | 0.03162 | 0.02980 | 0.03013 |
| 20 | 0.07374 | 0.02492 | 0.07175 | 0.02460 | 0.05315 | 0.02206 |
| 21 | 0.05216 | 0.02487 | 0.06503 | 0.02475 | 0.05265 | 0.02303 |
| 22 | 0.03797 | 0.02634 | 0.05389 | 0.02551 | 0.04878 | 0.02438 |
| 23 | 0.05600 | 0.02705 | 0.04833 | 0.02640 | 0.04511 | 0.02486 |
| 24 | 0.07043 | 0.04080 | 0.08087 | 0.03766 | 0.05687 | 0.03673 |
| 25 | 0.05688 | 0.03928 | 0.07117 | 0.03996 | 0.06221 | 0.03686 |
| 26 | 0.03085 | 0.02330 | 0.03030 | 0.02017 | 0.02893 | 0.01495 |
| 27 | 0.04549 | 0.01797 | 0.04557 | 0.01799 | 0.04479 | 0.01508 |
| 28 | 0.07780 | 0.04410 | 0.07528 | 0.04491 | 0.06335 | 0.04360 |
| 29 | 0.07750 | 0.04459 | 0.05613 | 0.04321 | 0.05173 | 0.04380 |
| 30 | 0.05205 | 0.04237 | 0.04386 | 0.03770 | 0.04007 | 0.01739 |
| 31 | 0.06236 | 0.02833 | 0.05826 | 0.02415 | 0.04872 | 0.01750 |
| 32 | 0.01072 | 0.03435 | 0.01079 | 0.04166 | 0.02839 | 0.00819 |
| 33 | 0.06160 | 0.01731 | 0.06227 | 0.01639 | 0.05234 | 0.00805 |
| $\bar{\sigma}_T^2$ | 0.0575 | 0.0322 | 0.0561 | 0.0354 | 0.0484 | 0.0281 |
| Median | 0.0573 | 0.0319 | 0.0552 | 0.0324 | 0.0487 | 0.0265 |

To further characterize testing results, we also performed pairwise comparisons of the variance estimates using the Wilcoxon sign-rank non-parametric test for paired samples. Fig. 8a shows the corresponding cumulative distribution function (CDF) estimates, while Fig. 8b shows the statistical test results. The figure shows that all estimates exhibit statistically significant differences (p < 0.05) with the exception of the relative comparison of the two variants of the energy based detection algorithm, either with or without preliminary filtering, indicating that the filtering does not play a major role.

**Figure 8.**
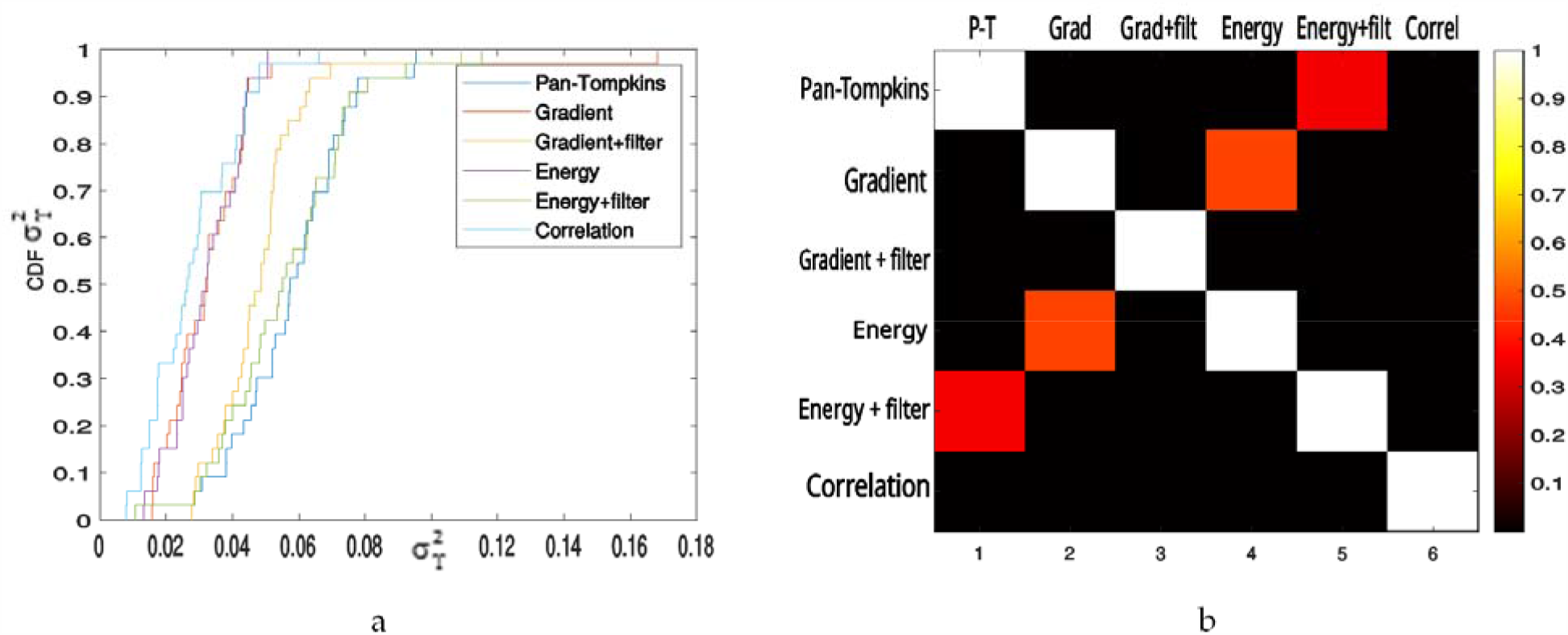
(a) Empirical CDFs of the total variance 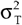 estimates obtained for the 33 tested long-term ECG records; (b) p-values by the Wilcoxon sign-rank non-parametric test for paired samples.

Altogether, our results indicate that the proposed gradient-based approach leads to the best positioning accuracy of single ECG cycles, indicated by the lowest total variance 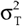 among tested approaches, outperforming the conventional Pan-Tompkins algorithm, while exhibiting comparable detection efficacy, indicated both by the analysis of average estimates and statistical test based comparisons. While the correlation based approach appears more accurate than the gradient-based one, it requires *a priori* knowledge of the particular QRS shape, while preliminary filtering does not help to improve the results.

### 4.2. Lane and band selection in the protein mass spectra analysis for the SDS-page fingerprinting

In the following, we show how similar strategy can be applied to the lane selection and band detection in the protein mass spectra analysis problem for the SDS-PAGE fingerprinting.

First, we consider the lane selection problem. A standard 60×85 mm electrophoresis gel photograph is averaged along the lanes to obtain the raw signal further used for the lane selection. In order to remove non-stationary background noise, a gliding average over 1/5 of the photograph size (corresponding to approximately two full lane widths) is subtracted, and a gradient estimate of the remaining fluctuating component is obtained and smoothed by averaging in a 10-pixel gliding window. The results of the automated lane selection algorithm are exemplified in Fig. 9, where panels (a)-(c) show examples of electrophoresis gel images, while panels (d)-(f) show the gradient based decision statistics ξ and the selection results based on the comparison of ξ with symmetric threshold levels ±θ. The choice of the threshold level is based on *a posteriori* statistics, i.e. several threshold levels are being tested, and that one that leads to the best results according to the optimization criteria is finally applied [29, 30]. Based on the assumption that lanes have more or less the same width, the minimum of the sum of the standard deviation of the lane width was chosen as the optimization criteria.

**Figure 9.**
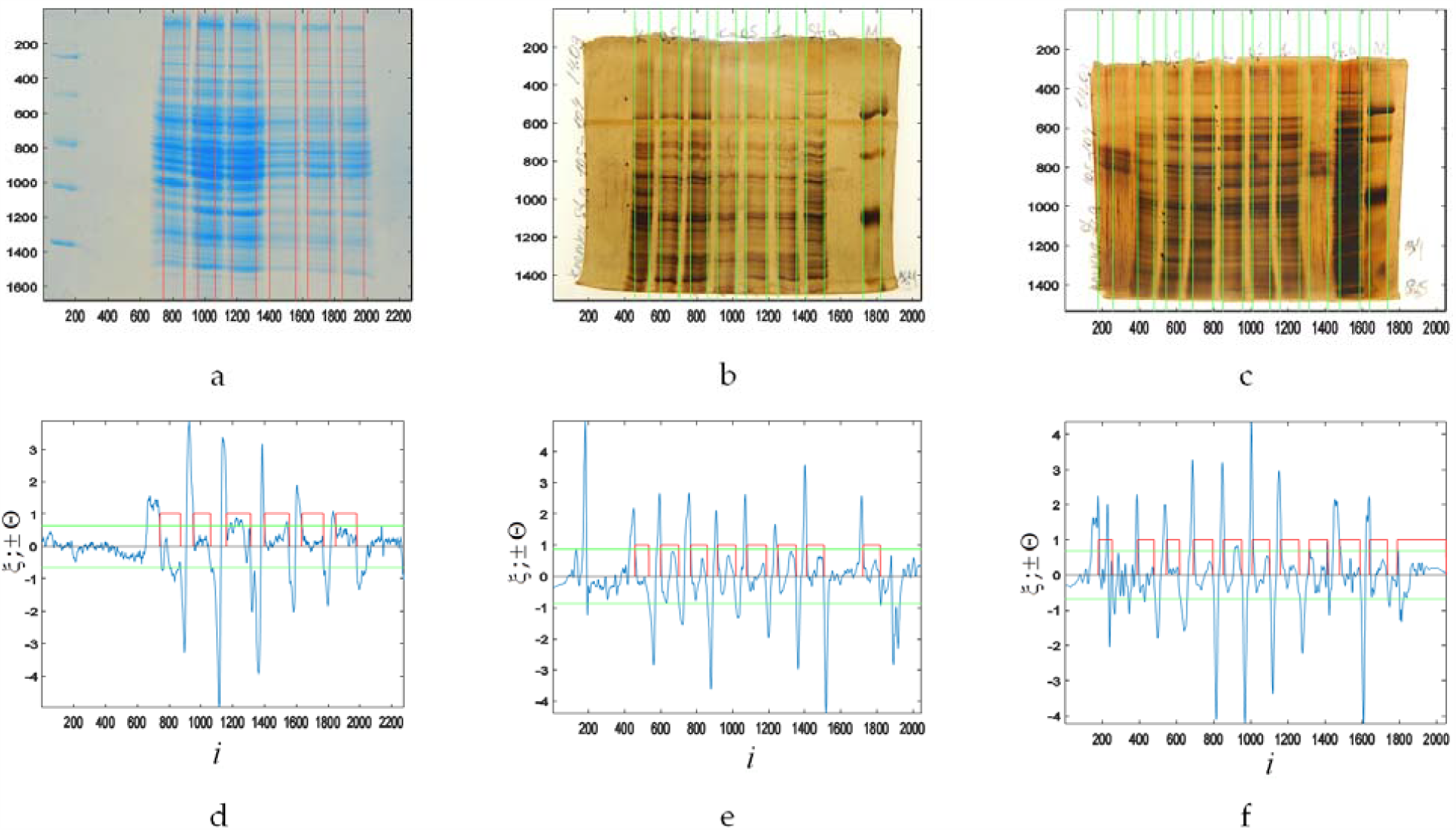
(a)-(c) Examples of electrophoresis gels photographs used in the SDS-page fingerprinting obtained from a recent study originally reported in [13], with highlighted results of the fully automated lane selection based on the proposed gradient based algorithm; (d)-(f) the gradient based statistics ξ used in the automated lane selection algorithm (blue curve) and the selection results (red bars) based on the comparison of ξ with threshold levels ±θ (green line), as a function of the respective image coordinate *i* (in pixels).

Finally, we applied the same gradient based approach to find individual protein bands in the selected electrophoresis gel lanes. For that, each selected electrophoresis gel lane has been averaged over its entire width w to obtain a one-dimensional signal with enhanced signal-to-noise ratio. The resulting signal was subjected to a simple detrending procedure by subtracting the baseline determined as the average of the lower quarter of its sample in a gliding window covering 1/10 of the lane length. Finally, a gradient estimate of the remaining fluctuating component was calculated as indicated above.

The results of the band selection are exemplified for two different electrophoresis gel lanes in Fig. 10, where panels (a) and (b) show the raw gel lane images, while panels (c) and (d) show the gradient based decision statistics ξ and the selection results based on the comparison of ξ with symmetric threshold levels ±θ. Finally, panels (d), (e) show the band selection results over the raw lane image.

**Figure 10.**
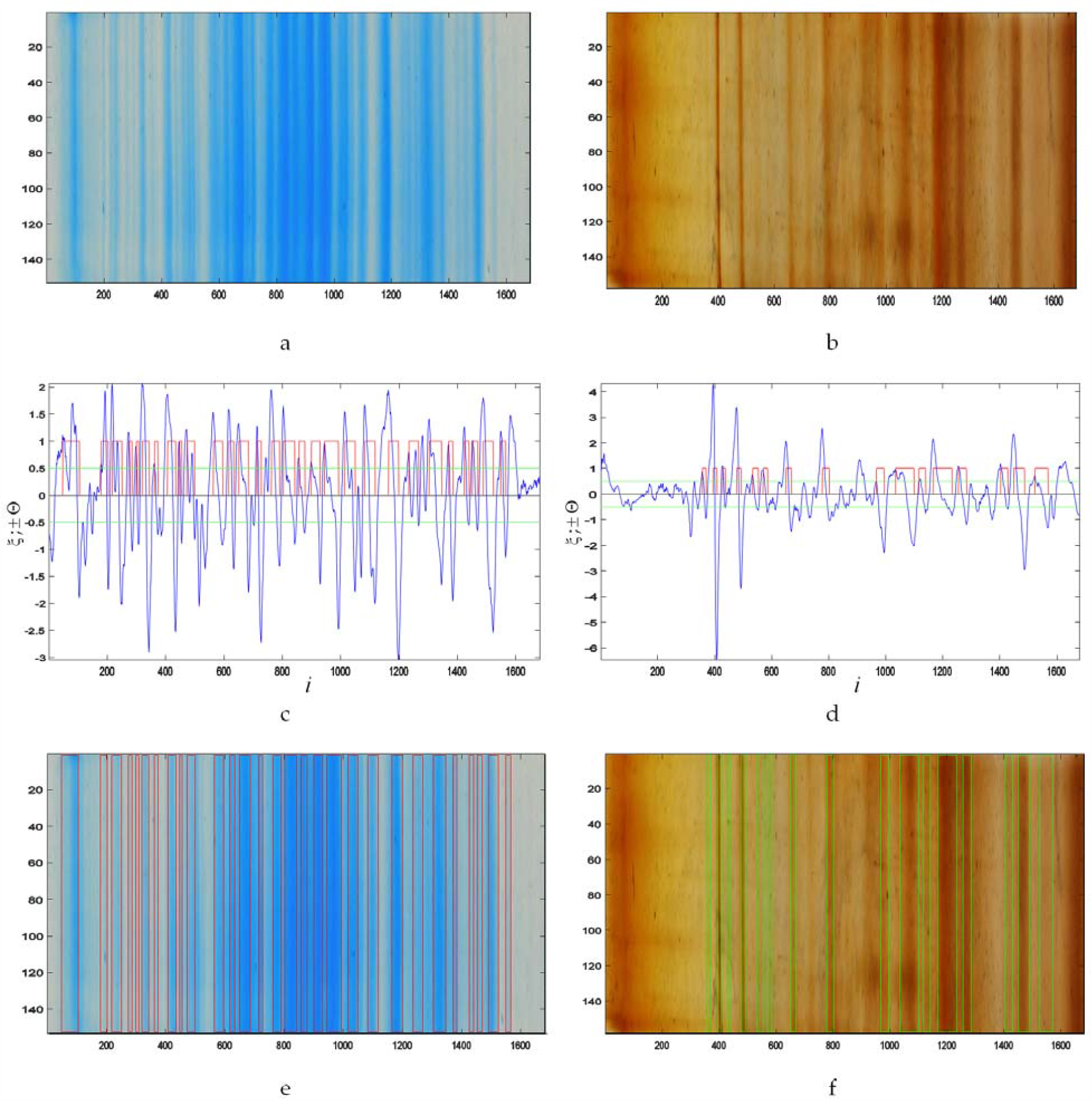
(a, b) Two representative sample lanes selected from electrophoresis gel images; (c, d) gradient based statistic ξ (blue curve) and threshold levels ±θ (green line) used in the band selection algorithm as a function of the respective image coordinate *i* (in pixels), selected bands (red bars); (e, f) results of the fully automated band selection over the raw gel images

### 4.3. Differentiation of protein mass spectra in the SDS-page fingerprinting

At last, we consider how the proposed transformations can be used to improve differentiation between nearly similar mass spectra by automated analysis of their electrophoresis gel lanes. In the majority of applications, nearly similar lanes that often differ by only one or few bands have to be compared, resulting in their similarity scores approaching one that is the worst case scenario in the differentiation problem.

Two representative examples shown in Fig. 11 support the above statement, indicating that the cosine similarity scores between the raw signals obtained from different electrophoresis gel lanes are around 0.97-0.99, making them hardly distinguishable between each other. Subtraction of the average from each lane, corresponding to the replacement of the cosine similarity score by the correlation coefficient, slightly improves the situation, while the centered signals remain far from being easily distinguishable from each other.

**Figure 11.**
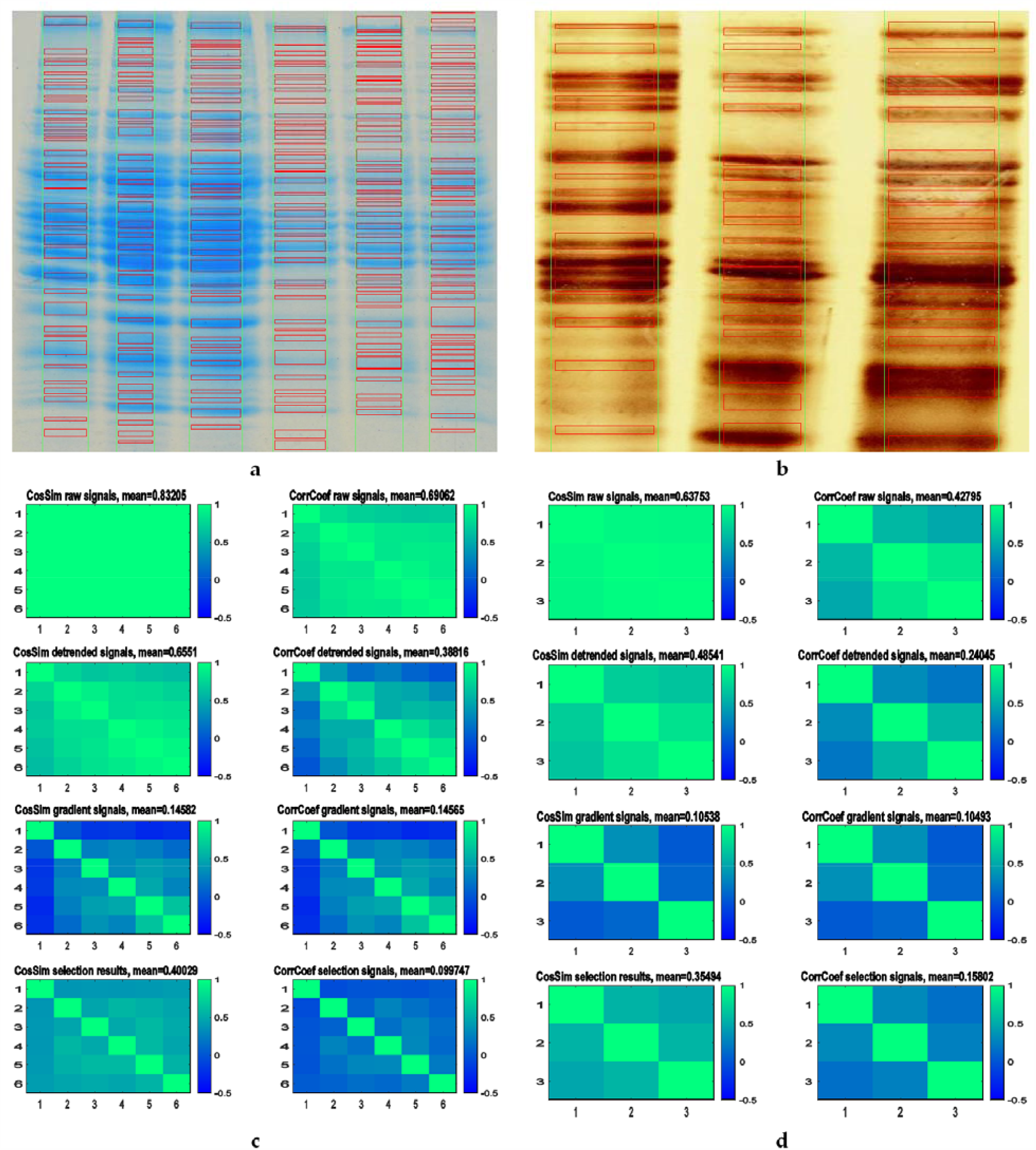
(a, b) Two sample electrophoresis gel images; (c, d) matrices of cosine similarity scores and cross-correlation coefficients of the raw, detrended, gradient signals, and binarized selection results. Matrix subtitles indicate average values obtained from all matrix entries with the exception of the main diagonal.

Further transformations following the proposed gradient based algorithm led to the reduction of both cross similarity metrics that eventually reached lowest similarity scores about 0.1-0.15, making distinguishing between such signals rather close to orthogonal signal ensemble scenario widely used in synthetic signal design [22]. In contrast, further binarization during the band selection step does not always lead to the improvement, as indicated by the corresponding similarity scores. It is therefore questionable, whether the last step is always required, as it does not allow for any intermediate scenarios, like reduced intensity lanes exemplified in the lower left corner of Fig. 8b. Depending on particular problem setting, either comparison between binary selection results, or comparison directly between gradient based signal transformations could be advisable.

Finally, depending on the problem setting, there are two scenarios of the lane differentiation problem. The simplest scenario is distinguishing between the two lanes, which usually involves running a statistical test based on some similarity score, which provides with a confidence probability that the signals obtained from the two lanes are identical. The second scenario is finding the best match to the tested sample in a series of reference lanes, which is resolved by finding the maximum similarity score. Finally, there is a third possible scenario, when one expects that one of the reference lanes might correspond to the tested sample, although there is no certainty that there is a true matching lane available in the reference ensemble. It this latter scenario, based on the assumption that the tested and the reference lane signals *x*_*i*_ *= s*_*i*_ *+ n*_*i*_ are assembled by identical signal samples *s*_*i*_ and different noise samples *n*_*i*_ can be used to estimate the ROC curves like in Fig. 1 and estimate the potential error from their overlap. In the first approximation, one can estimate the confidence interval for the self-similarity score as a certain quintile of the similarity score distribution that, in turn, can be approximately calculated based on Eq. (5) obtained by analytical treatment. By setting a threshold at the 5% quintile of the respective distribution, one can assume that the those lanes exceeding the threshold value are identical with 95% confidence probability, since their similarity scores appear within the 95% confidence interval for the self-similarity score. Any signal transformation like taking the gradient requires recalculation of this threshold value following Eq. (5).

Figure 12 shows a prominent example of multiple lanes comparison, where a matching lane is marked by green, and other lanes are marked by red. The figure shows that in both cases, whether the raw signals or their gradient transformations are being compared, the highest similarity score corresponds to the matching lane. However, in the case of the raw signals comparison, about one half of all similarity scores exceed the threshold, indicating that they appear within the 95% confidence interval for the self-similarity score estimate. In contrast, in the case of the gradient based signal transformations comparison, only one single similarity score corresponding to the matching lane exceeds the threshold, indicating that differences between the tested lane signal and other reference signals are well below the decision threshold.

**Figure 12.**
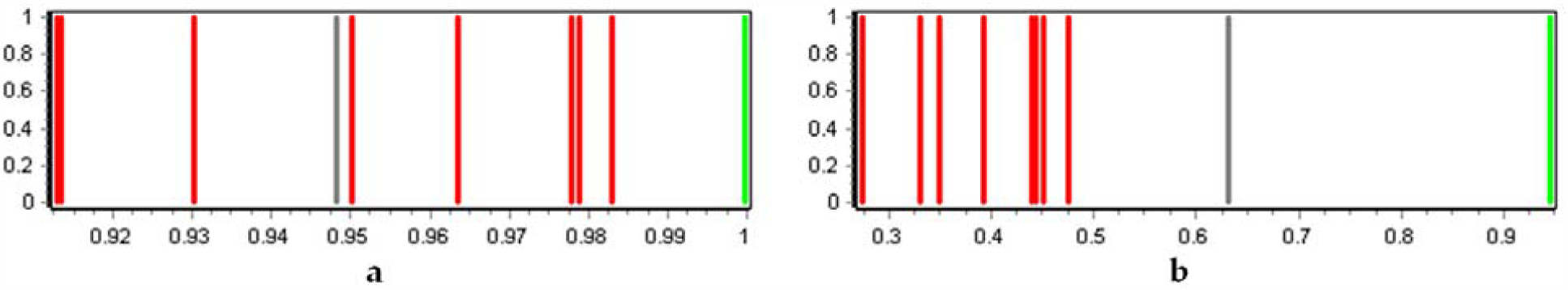
Similarity scores based on (a) raw signal and (b) their gradient based transformations comparison in a signal differentiation problem for an electrophoresis gel image example.

## 5. Conclusions

To summarize, we have proposed a simple and universal approach to the online detection of events represented by abrupt bursts in long-term observational data series based on a simple gradient-based nonlinear signal transformation obtained as a product of the signal and its derivative. We provided explicit analytical expressions characterizing the efficacy of the proposed gradient-based approach in comparison with the conventional solutions optimized for particular theoretical scenarios and widely utilized in various signal analysis applications. We have shown that an improved online detection of any significant bursts in the observational data series irrespective of their particular shapes could be achieved. Moreover, we show explicitly, that the gradient-based approach outperforms the conventional Pan-Tompkins algorithm in its original formulation in the exact positioning of single ECG cycles, while exhibiting comparable detection effectiveness. Finally, we show that our approach is also applicable for the automated detection of individual bands and comparative analysis of lanes in electrophoretic gel images. As an outlook, we think that further improvements might be achieved by taking into account memory effects in the heartbeat sequence records [31, 32], which could itself act as an additional diagnostic indicator [33]. Also similar approach might be helpful for the online analysis of certain secondary physiological indices that could be derived online from the recorded physiological signals, such as the baroreflex sensitivity [34].

Since detection of events represented by abrupt bursts irrespectively to their particular shapes is of immense importance for the improvement of biomedical signal analysis systems, we believe that our findings could be useful for the design of such systems, as well as for the use in molecular diagnostic systems exploiting the biomolecules mass spectra analysis such as the restriction fragment length polymorphism (RFLP) and the variable number tandem repeats (VNTR). A simple software tool for the semi-automated electrophoretic gel image analysis based on the proposed gradient based methodology for the lane selection and band detection, as well as signal comparison in different lanes, is freely available online [35] and can be used for the signal analysis in the SDS-PAGE fingerprinting or similar experimental data processing.

## Acknowledgements

We like to acknowledge the financial support of this work by the Ministry of Science and Higher Education of the Russian Federation under assignment No. 0788-2020-0002.

